# mtDNA-Network: a web tool to explore mitochondrial variant profiles in complex diseases

**DOI:** 10.64898/2026.01.15.699732

**Authors:** Leticia Cota Cavaleiro de Macêdo, Gustavo Barra-Matos, Marcella Vitória Belém-Souza, Camila Sinimbú Forte, Helber Gonzales Almeida Palheta, Ândrea Ribeiro-dos-Santos, Giovanna Chaves Cavalcante, Gilderlanio Santana-de-Araújo

## Abstract

The mitochondrial genome (mtDNA) provides valuable insights into human evolution, population diversity, and disease etiology. Here, we present the mtDNA-network (https://apps.lghm.ufpa.br/mtdna/), an integrative bioinformatics database and tool for the visualization and analysis of mitochondrial variants (single-nucleotide variants and insertions/deletions). The mtDNA-network was upgraded to enable investigation of mtDNA in admixed Brazilian individuals with substantial contributions from uniparental Indigenous and African ancestries. We implement a bioinformatics pipeline to harmonize variant calling across 339 mtDNA samples. The dataset supports general genetic population analysis and variant mapping for complex diseases, including Parkinson’s disease (104 cases and 75 controls), leprosy (33 cases and 37 controls), and somatic gastric cancer (40 cases and 50 controls). The mtDNA-network tool features an intuitive interface and analytical dashboards for transitions, transversions, heteroplasmy, and variant-disease networks, and serves as a strategic resource for advancing research in population genetics and precision medicine in underrepresented populations. We reinforce the critical need to expand non-European genomic representation in global databases to promote more equitable genomic diversity studies and clinically relevant discoveries.

## 1. Introduction

Mitochondria are essential cytoplasmic organelles that perform essential molecular functions, including oxidative phosphorylation (OXPHOS). Due to their evolutionary origin, these organelles possess their own genetic material, known as mitochondrial DNA (mtDNA), which is circular, double-stranded, and encodes key components of the electron transport chain (ETC) as well as ribosomal and transfer RNAs. Given its proximity to reactive oxygen species (ROS) generated during ATP synthesis and its limited repair mechanisms, mtDNA is particularly vulnerable to mutations, deletions, depletion, and copy number variation that can disrupt mitochondrial function [1].

Mitochondrial dysfunctions have been linked to numerous human diseases. In cancer, mtDNA mutations can drive tumorigenesis by altering metabolism, increasing ROS production, and promoting metastasis [2–4]. In neurodegenerative disorders such as Alzheimer’s disease and Parkinson’s disease, impaired mitochondrial function may contribute to neuronal loss through energy failure and oxidative stress [5,6]. Moreover, in infectious diseases such as leprosy, pathogens control host mitochondria to enhance survival and replication [7,8]. For population genomics studies, significant challenges persist in building comprehensive and equitable genetic databases for underrepresented non-European populations [9]. Most platforms lack computational frameworks that link genetic variation, evolutionary patterns, and disease associations, thus limiting insights into complex genotype-phenotype relationships, particularly in admixed populations [9]. In contrast to nuclear data, mtDNA information remains fragmented and poorly annotated, delaying large-scale analyses and accurate haplogroup classification [10]. This limitation is particularly critical for tracing maternal lineages in highly admixed populations, such as Brazil. The conception of curated, high-resolution mtDNA databases and tools that characterize diverse populations is essential to advance clinical and population genetics.

Here, we present the new version of mtDNA-network, an interactive bioinformatics web platform that integrates genomic, genealogical, and clinical data to visualize relationships between mtDNA variants and complex diseases. The mtDNA-network, which features dynamic dashboards, interactive tables, and network modeling, enhances research on the genetic diversity and disease associations, with a particular focus on underrepresented populations enriched with Brazilian Indigenous and African ancestry.

## 2. The mtDNA-network

The mtDNA-network (https://apps.lghm.ufpa.br/mtdna/) is an integrative bioinformatics platform that connects genomic, genealogical, and clinical information through interactive graphical representations. The tool enables researchers across fields, including geneticists, anthropologists, and physicians, to explore the relationships between mtDNA variants and complex diseases, thereby advancing understanding of their etiology, particularly in underrepresented populations characterized by complex admixture of Indigenous and African ancestries. Compared with the first version [11], we have substantially expanded the mtDNA-network to incorporate whole mitochondrial genome sequencing data for population genetics and disease association analyses. The updated resource now includes data from individuals affected by gastric cancer, Parkinson’s disease, and leprosy, allowing comparative studies in various pathological contexts. The platform integrates scalable, high-performance genetic analytics modules that are user-friendly. New features include interactive tables, dynamic dashboards, and advanced network modeling tools that facilitate the investigation of genetic diversity and its biomedical implications, particularly within underrepresented populations characterized by a complex admixture of deep Indigenous and African ancestry. The interactive web interface provides modules for exploring variant dashboards, haplogroup visualization, and network modeling and visualization (Figure 1A). The tool also includes comprehensive online documentation to support users, developers, and system maintenance, thereby promoting reproducibility and community-driven development for future expansion. At its core, the mtDNA-network implements a variant detection pipeline that encompasses modules for identifying single-nucleotide variants (SNVs) and insertions/deletions (INDELs). The pipeline is available on GitHub (http://www.github.com/gilderlanio/mitogenome-variant-calling) and Zenodo (https://zenodo.org/records/16617240) and provides a reproducible and scalable workflow for mitochondrial genome analysis (Figure 1B).

**Figure 1.**
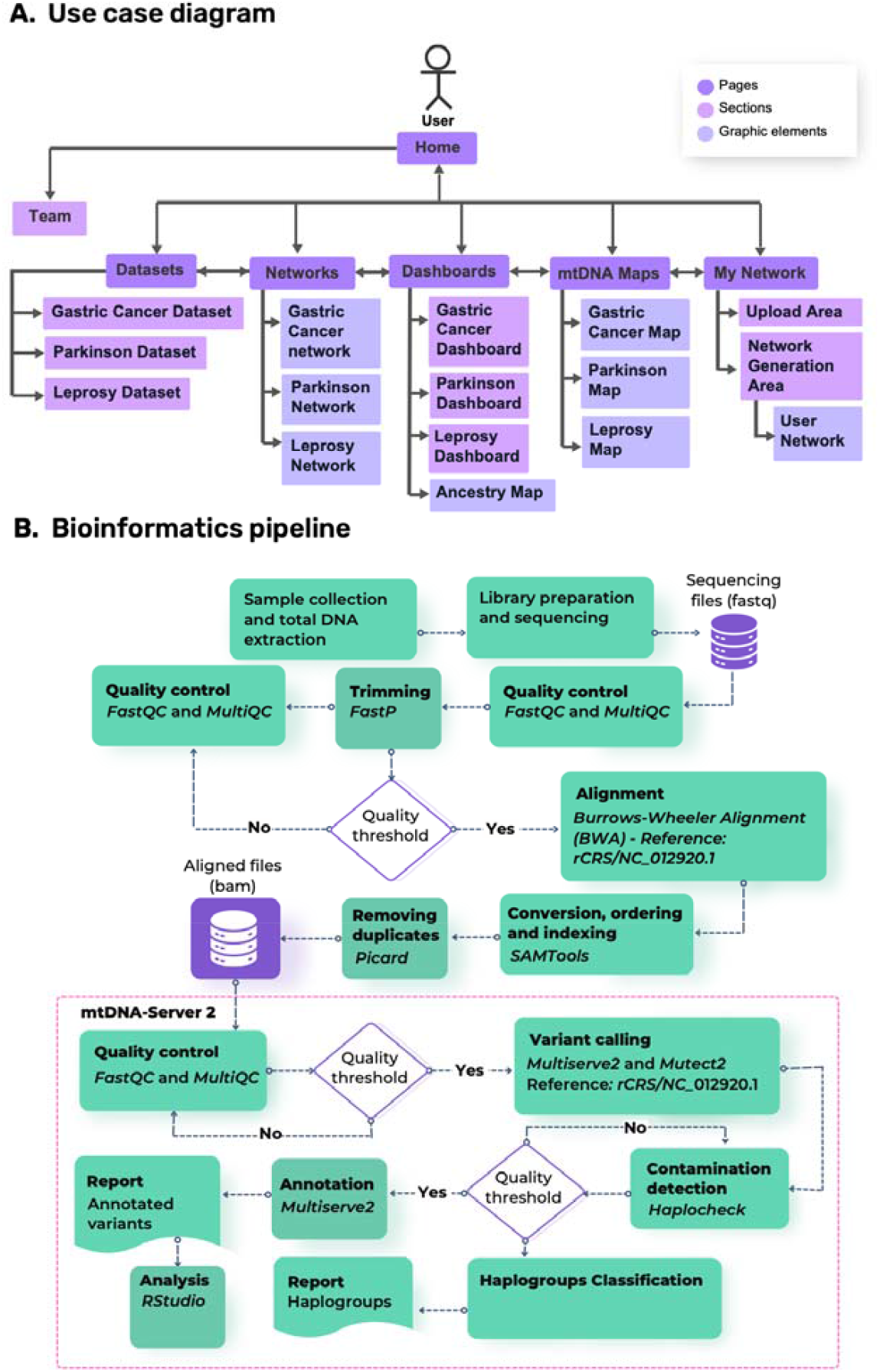
Overview of the mtDNA-network platform and pipeline. (A) Use case diagram for interacting with the mtDNA-network platform. Each module allows for variant exploration, haplogroup visualization, and network modeling. (B) The variant-calling pipeline to identify SNVs and INDELs in mitochondrial genomes. The pipeline is available on GitHub (http://github.com/gilderlanio/mitogenome-variant-calling) or in Zotero.

## 3. mtDNA sequencing data

The first version of the tool was designed to analyze the profiles of genetic variants in gastric cancer compared with 36 other cancer types [11]. In this version, we included two additional mitogenome sequencing databases conducted by our research group, deposited at the European Nucleotide Archive under the accession numbers PRJEB59275 and PRJEB74357, comprising 343 samples obtained from Brazilian individuals, corresponding to the following studies:

- For gastric cancer, mitochondrial genome data are available from Cavalcante et al. [12], who compared mutation profiles in gastric tissue between healthy controls (n = 50) and gastric cancer patients (n = 40). This tool incorporates publicly available data from multiple sources, including The Cancer Mitochondrial Atlas (TCMA) [4].
- For Parkinson’s Disease (PD): The mitogenome from Matos et al. (2024) comprises 96 individuals, to which 87 additional individuals were added, divided into case and control groups. The case group comprises 106 patients with idiopathic PD: 66 without levodopa-induced dyskinesia (NLID) and 40 with levodopa-induced dyskinesia (LID) [6]. The control group consisted of 77 individuals without a diagnosis of PD or other neurodegenerative disorders.
- For leprosy: Mitogenome data from leprosy patients (case group, n=33) and healthy household contacts (control group, n=37) were integrated from [7,8]. The case group was subdivided into three subgroups based on leprosy type: borderline lepromatous (BL) (n=12), lepromatous (LL) (n=11), and borderline tuberculoid (BT) (n=10).

## 4. mtDNA bioinformatics pipeline

The whole-genome workflow is drawn in Figure 1B, including pre-processing of .fastq and variant calling. Initial quality assessment of raw sequencing data was performed using FastQC (v0.12.1) and MultiQC (v1.19) to evaluate read quality before and after processing [12,13]. Preprocessing of mitochondrial DNA sequences involved adapter trimming, removal of low-quality bases (Phred score < Q30), and exclusion of reads shorter than 36 nucleotides using FastP (v0.23.4) [14]. The trimming process targeted low-quality bases (Q < 30) at the read end, followed by stringent filtering to discard reads and systematic adapter removal to minimize technical artifacts. Processed reads were aligned to the Revised Cambridge Reference Sequence (rCRS; NC_012920.1) using the Burrows-Wheeler Aligner (BWA v0.7) [15]. The resulting SAM files were converted to BAM format, sorted, and indexed using SAMtools (v1.15.1) [16]. To reduce PCR bias, duplicate reads were marked and removed using Picard (v2.27.5) [17]. Sample contamination was rigorously assessed with Haplocheck (v1.3.3), and samples exceeding the 10% contamination threshold were excluded from subsequent analyses [18]. High-confidence variants, including SNVs and INDELs, were identified using mutserve and classified as transitions (TS) or transversions (TV). To ensure reliability, only variants with mean read depth spanning the mitochondrial genome and heteroplasmy levels between 5% and 95% were retained. Samples with contamination >10% were excluded. Filters based on read depth and heteroplasmy were applied throughout the analysis.

## 5. Software Architecture

The mtDNA-Network is a web-based computational platform built with the Flask microframework that provides an interactive, integrated environment for the analysis and visualization of mitochondrial DNA variant data. The system was structured with a modular architecture that enhances scalability, maintainability, and efficient communication between client-side and server-side components. As depicted in Figure 2, the root directory contains the primary executable script, responsible for initializing the Flask application and importing its associated modules, as well as two principal directories: templates, which store the HTML files defining the user interface, and static, which hosts static resources, including images, JavaScript files, CSS stylesheets, and data files used in tabular and graphical representations.

**Figure 2.**
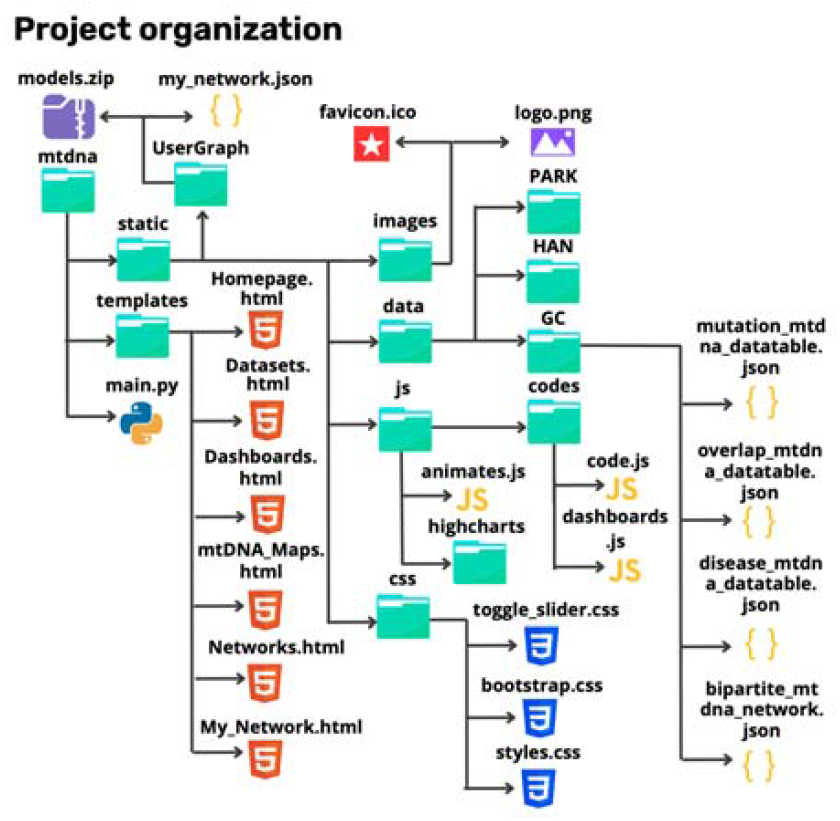
Overview of the software architecture, illustrating the organization and relationships among the main directories and files that constitute the system.

The platform integrates specialized libraries to support advanced data visualization and interactive user engagement. Tabular data was displayed using the DataTables plugin (jQuery), which enables dynamic filtering, pagination, and sorting of large-scale biological datasets. Network-based visualizations were implemented using D3.js, allowing the interactive rendering of complex network topologies and facilitating real-time exploration of relationships among mitochondrial haplotypes or sequence variants.

In addition to network analysis, mtDNA-Network incorporates multiple graphical frameworks to provide complementary analytical perspectives. Interactive dashboards were generated using HighCharts.js, whereas UpSet plots were constructed with Vega-Lite, a declarative visualization grammar conceptually analogous to ggplot2 in R. Circular representations of the mitochondrial genome were produced with CGView.js, offering accurate and intuitive visualizations of genomic organization and functional regions.

Data communication between the client interface and the backend were handled via Flask-CORS APIs and POST requests, enabling the dynamic creation, modification, and deletion of JSON files required for user-specific network generation. This architecture supports responsive updates and efficient data management while maintaining lightweight server-side processing. The web interface combines custom CSS with Bootstrap, Animate.css, and Tilt.js to deliver a responsive, visually cohesive layout that improves usability, accessibility, and cross-device compatibility.

## 6. Visualization features

The mtDNA-Network platform integrates an extensive suite of visualization tools specifically designed for the interactive exploration of mtDNA variants, including SNVs and INDELs. These features enable users to analyze, compare, and interpret genetic alterations across a range of pathological conditions and at the population level. The following subsections describe in detail the main visualization components, highlighting their functionalities and applications for genetic and bioinformatic analysis.

### 6.1. Dataset view

The Datasets view serves as the basis for raw data exploration within the platform. Provides dynamic, interactive tables that enable users to systematically navigate large-scale variant data and compare variants. For each disease, a Dataset view was created with structured tables that summarize genetic variants, associated genes, and relevant metadata for each study cohort. For the cancer data set view, the platform displays all identified variants, including somatic mutations, along with their genomic positions and associated genes. Each cancer subtype was characterized by a distinct mutational profile, enabling comparative analysis across tumor types. The degree of similarity between different cancers was quantified using the Jaccard index, which measures the proportion of shared variants between two datasets relative to the total number of unique variants. This quantitative approach allows researchers to assess the overlap of mutational spectra across cancers, supporting the identification of potential common mitochondrial alterations or pathways involved in oncogenesis. For the leprosy dataset, the platform provides a detailed catalog of all mitochondrial SNVs detected in the analyzed samples. Each variant was annotated with its predicted functional consequence, such as a nonsynonymous substitution, a synonymous change, or a mutation in non-coding regions. Additional biological and clinical metadata were integrated into the visualization, including heteroplasmy levels, patient ancestry, and clinical subtype (LL, BL, and BT). In the Parkinson’s disease dataset, the system catalogs all variants detected SNVs and INDELs for each individual in the control and disease groups. Each entry includes the affected gene, the variant classification (e.g., synonymous, nonsynonymous, or frameshift), sequencing coverage depth, and heteroplasmy percentage. The Datasets view for each disease incorporates several advanced functionalities that enhance data interaction and usability. Users can apply filters based on variant type, gene, or disease association, reorder columns, and paginate large datasets without compromising performance. In addition, a full-text search feature enables rapid retrieval of specific variants or genes, and all tables can be exported in standard formats for further bioinformatic or statistical analysis using external software.

### 6.2. Analytical dashboards view

Complementing the tabular datasets, the Dashboards view module provides a set of highly interactive analytical panels to summarize and visualize genetic variants. These dashboards transform raw variant data into intuitive graphical summaries that reveal patterns, distributions of mtDNA variants, and relationships for case-control studies. Statistical summaries were graphically presented, including comparisons of variant frequencies between cohorts; transition-to-transversion (TS/TV) ratios by disease group; distributions of heteroplasmy levels across genes; and shared variant patterns between diseases, expressed through overlap and similarity graphs. The dashboards also display Jaccard indices (gastric cancer ensemble) comparing variant sets, variant counts per gene, and the ancestry composition of individual samples, allowing researchers to correlate genetic diversity with population structure. Each graphical element is fully interactive. Users can hover over data points to reveal contextual information through dynamic tooltips, zoom in on specific data regions, and apply filters directly within the visualization. All charts can be exported in multiple formats suitable for publications and reports. Some visualizations, such as UpSet plots, include a dedicated view for inspecting and exporting the underlying code, thereby supporting transparency and reproducibility in data analysis workflows. Figure 3 shows some of the graphical charts from the PD session, such as the distributions of transitions and transversions (Figure 3A and 3 B), the mtDNA circle plot with the distribution of variants (Figure 3C), a donut chart with the number of INDELs at each locus (Figure 3D), a bar graph with the number of individuals per haplogroup (Figure 3E), an upset with the variants shared between the case and control groups (Figure 3F), and a boxplot with the relationship between sex-by-age between individuals (Figure 3G).

**Figure 3.**
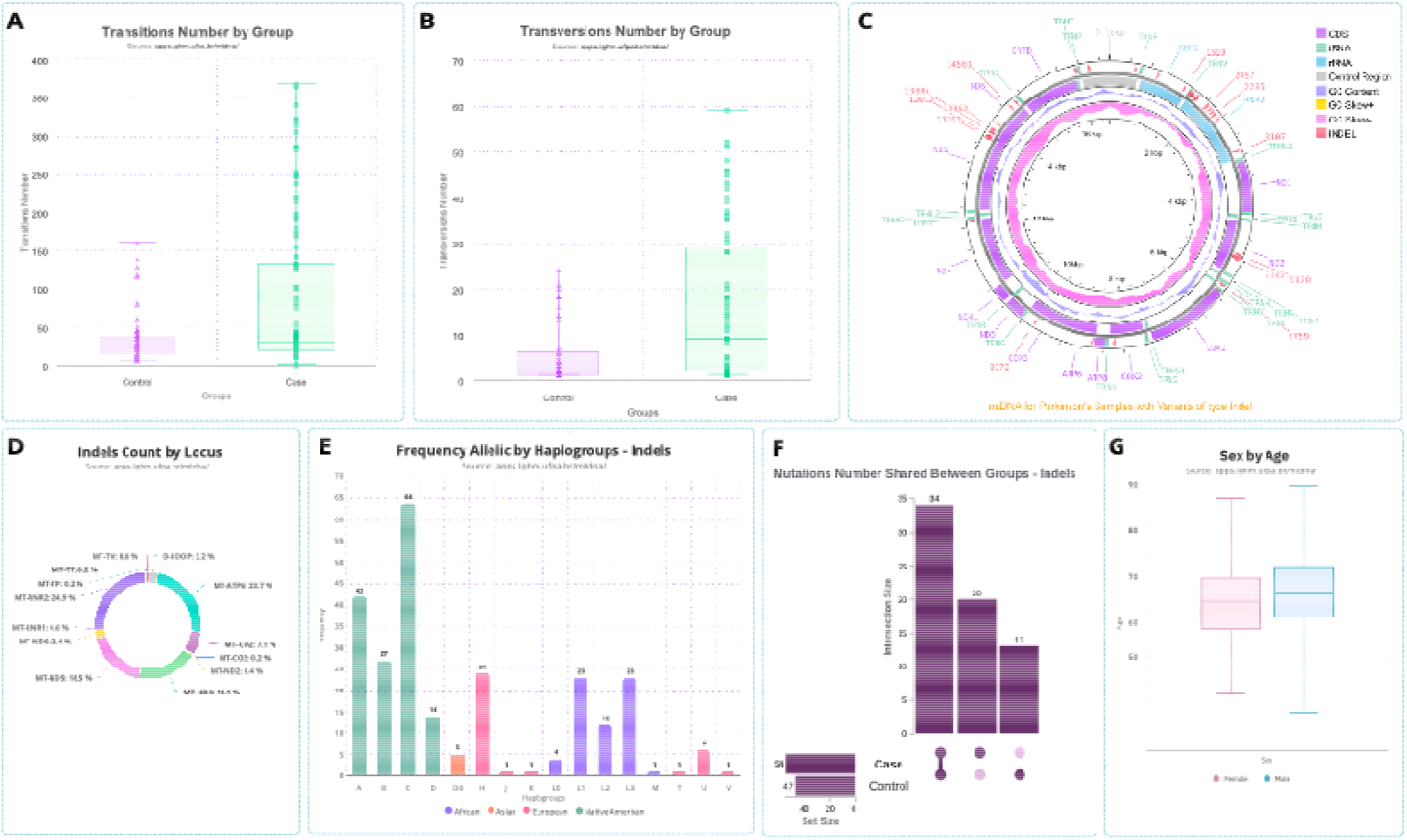
Example visualizations from the Parkinson’s disease section of the mtDNA maps and dashboards. These include distributions of individuals by sex, transition/transversion ratios by group, INDEL counts by gene, allele frequencies by haplogroup, and shared variants between cohorts. (A) Frequency of transition mutations observed in Parkinson’s disease cases. (B) Frequency of transversion mutations observed in Parkinson’s disease cases. (C) Circos plot showing the genomic distribution of mtDNA variants across loci. (D) Number of INDELs variants per mitochondrial DNA locus. (E) Distribution of mitochondrial haplogroups represented in the mtDNA network. (F) Overlap of mtDNA variants between Parkinson’s disease cases and controls. (G) Sex distribution of study participants is shown as a boxplot.

### 6.3. Macrohaplogroup overview

The mtDNA-network analysis encompasses the germline mitogenomes of 249 individuals from the North region of Brazil, providing insights into the maternal genetic landscape of this population. Using HaploGrep3, sixteen distinct macrohaplogroups were identified and classified with high confidence (posterior probability > 0.89) into four major ancestral origins: Indigenous (A, B, C, and D), European (H, J, K, T, U, and V), African (L0, L1, L2, L3, and M), and Asian (D4) (Figure 3E). The presence of these diverse haplogroups reflects the region’s complex demographic history, shaped by multiple waves of migration, colonization, and admixture over centuries. Among Indigenous haplogroups, haplogroup C emerged as the most prevalent, consistent with previous genetic studies characterizing Indigenous populations in northern Brazil. This predominance further supports historical and anthropological evidence that indicates the long-standing settlement and continuity of specific maternal lineages in the Amazon basin. The distribution of macrohaplogroups observed in this cohort not only corroborates known patterns of Native American ancestry in the region but also provides a refined understanding of the matrilineal contributions to the genetic structure of contemporary admixed populations in northern Brazil.

### 6.4. mtDNA Maps view

For genomic and phylogenetic interpretation, the mtDNA maps module provides high-resolution, interactive representations of the mitochondrial genome. Variants were aligned to the NCBI reference sequence (NC_012920.1), enabling users to visualize their distribution along the circular mitochondrial genome or in a linear representation. These maps were annotated with genomic features, including protein-coding genes, rRNA and tRNA loci, and other functional regions, all presented with nucleotide-level precision (Figure 3C). The platform supports smooth zooming and panning, allowing exploration of the whole mtDNA. Users can also download these visualizations in multiple formats for inclusion in manuscripts, reports, or presentations.

### 6.5. Networks and Custom Networks view

To reveal relationships between genetic variants and phenotypic traits, the Network visualization module implements bipartite graph representations, with nodes corresponding to genetic variants or phenotypes, following the approach of Araújo et al. [19,20]. This design enables users to explore associations and shared features across groups, facilitating the detection of patterns that are not readily apparent in traditional tabular data. The Parkinson’s disease network (Figures 4A and 4B) focuses on INDELs and SNPs associated with the control, dyskinesia (LI), and non-dyskinesia (NLID) groups, enabling visual differentiation of mutation patterns between clinical subgroups. In the gastric cancer network (Figure 2C), data from this study are integrated with entries from the TCMA database, with cross-dataset relationships highlighted by color-coded nodes (pink for TCMA-derived variants). The leprosy network (Figure 4D) maps heteroplasmic SNPs to clinical subtypes (LL, BL, and BT), providing insight into genotype–phenotype associations and potential mitochondrial contributions to disease progression.

**Figure 4.**
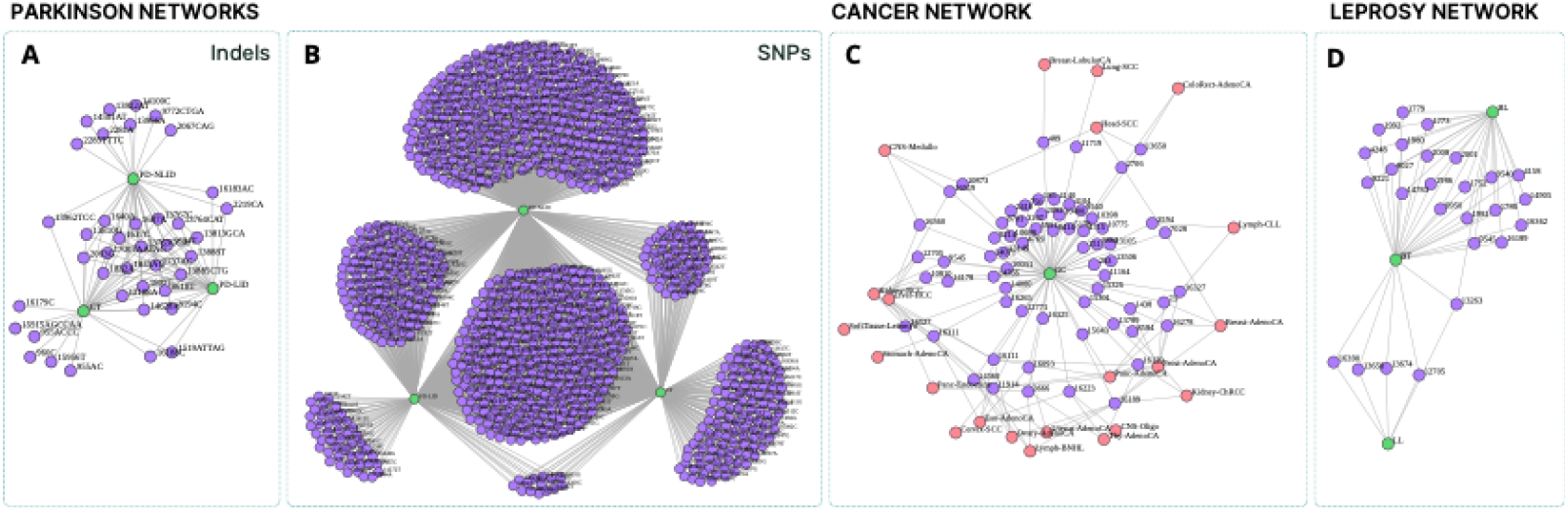
Bipartite networks for the three case studies integrated into the mtDNA network. Each network depicts associations between phenotypes and mitochondrial variants, with nodes representing either SNVs or INDELs mapped to each phenotype.

All proposed interactive network visualizations are fully dynamic tools that support filtering, zooming, node highlighting, and network reorganization. Through these capabilities, researchers can generate hypotheses, identify co-occurring variants, and stratify patients according to shared mitochondrial variant profiles. A compelling feature is the Custom Network Generator, which allows users to import their own datasets and construct personalized bipartite networks using the same visualization framework. Researchers can upload .csv files containing three essential columns: node identifiers, optional node colors, and connections between nodes. Links can be defined either one per row or separated by the | character, as shown in Table 1. Once uploaded, these networks retain all interactive features of the pre-existing ones, enabling full customization of visual parameters and layout. Figure 5 shows the upload area, where the user imports the .csv file, and the graphical generation area, where the network is rendered from the imported .csv file. This functionality facilitates the integration of external datasets, enabling cross-study comparisons, hypothesis validation, and the extension of the platform for collaborative and comparative research projects in mitochondrial genetics.

**Table 1.** Example of a network imported into the network module, where each row represents a node, its assigned color, and the nodes to which it is connected.

| nodes_id | nodes_color | connected_nodes |
| --- | --- | --- |
| A | green | B |
| B | purple | Z |
| C | purple | A \| B |
| ⋮ | ⋮ | ⋮ |
| Z | pink | A \| B \| C |

**Figure 5.**
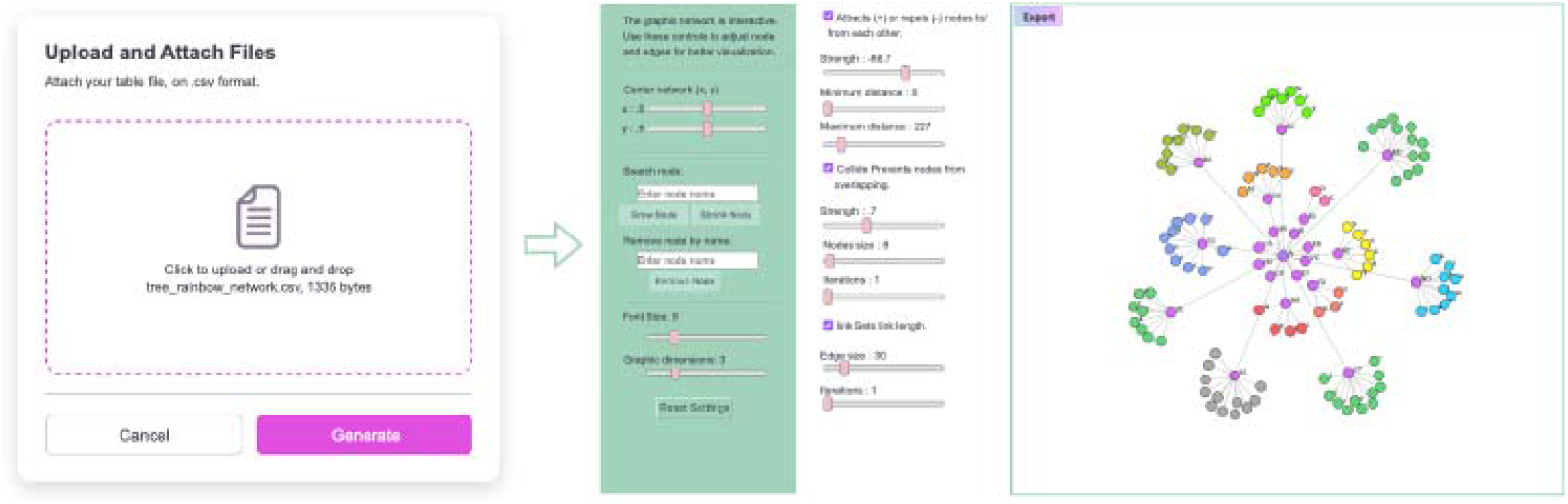
The mtDNA-network interface for generating custom networks. The left panel shows the My Network upload area, where users import a .csv table to create an mtDNA network. Upon clicking Generate, the network is automatically rendered in the panel on the right, where it can be interactively customized and exported.

## 7. Discussion

The mtDNA-Network web tool is a computational and bioinformatics resource designed to address a critical gap in mitochondrial genomics studies. The mtDNA plays a crucial role in cellular bioenergetics and has been increasingly implicated in the etiology of a wide range of complex diseases, including neurodegenerative disorders, metabolic syndromes, and various cancers [10,5,21].

A comprehensive analysis of mtDNA variants, particularly within underrepresented populations such as the Brazilian cohort, remains challenging due to the scarcity of dedicated bioinformatics tools and data for supporting accurate genetic diversity studies [10]. The mtDNA-network addresses this gap by providing a robust, integrated platform for exploring mtDNA variants. Notably, we are contributing to studies of populations with significant uniparental genomic contributions from Indigenous and African populations, and to the generation of biologically meaningful hypotheses regarding genotype-phenotype associations.

The current mtDNA-Network constitutes a substantial advancement in introducing new analytical and visualization features. Notably, the enhanced pipeline supports the detection of mitochondrial mutations, SNPs, and INDELs derived from whole mitogenome sequencing. Furthermore, the integrated pipeline tool enables haplogroup classification, facilitating analysis of mtDNA lineages. This is particularly valuable for population genetics studies, as it enables delineation of mtDNA diversity across ancestral backgrounds, including African, European, Asian, and Indigenous lineages.

Importantly, the mtDNA-Network is the first computational tool to offer an integrated analysis framework explicitly tailored to the Brazilian population, linking mtDNA variants to disease associations [5]. The unique genetic landscape of the Brazilian population, shaped by the admixture of European, African, and Indigenous peoples, has historically been underrepresented in global genomic databases and tools.

This innovative feature makes a significant contribution to medical genomics and epidemiology, enabling researchers to systematically and scalably explore the pathogenic potential of population-specific variants. Through dynamic graphical interfaces and intuitive data representations, users can efficiently navigate complex datasets, streamline the interpretation of results, and refine biological hypotheses.

Overall, the mtDNA-Network architecture emphasizes modularity, interactivity, and performance, ensuring a fluid integration between visualization, analysis, and user interaction. By combining robust web technologies with modern data visualization frameworks, the platform provides an efficient and user-friendly environment for exploring mitochondrial datasets, thereby facilitating the interpretation and discovery of biologically relevant patterns.

## 8. Conclusion

The mtDNA-Network tool offers a powerful, innovative platform to advance genomic research on underrepresented populations, particularly in relation to complex diseases, such as Parkinson’s disease, leprosy, and cancer. By combining robust analytical capabilities with an intuitive, user-friendly interface, the tool enables more effective interpretation of complex genetic data, bridging the gap between raw genomic information and clinically relevant insights. Thus, the mtDNA-Network platform stands out as a critical resource for researchers exploring mitochondrial genomics across populations and identifying novel associations between mtDNA variants and disease phenotypes.

## CRediT authorship contribution statement

L. C. Cavaleiro de Macêdo: Development of the new version of the software and its periodic updates. G. B. Matos: Development of the bioinformatics pipeline and data pre-processing. F. G. de Souza: Manuscript revision. C. S. Forte: Tool validation. M. V. B. Souza: Tool validation. C. S. Silva: Manuscript revision. H. G. A. Palheta: Manuscript revision and guidance for including the project on the laboratory server. B. L. Santos-Lobato: Raw data provider. Â. Ribeiro-dos-Santos: Raw data provider. G. C. Cavalcante: Conceptualization, Writing – review editing. G. S. de Araújo: Supervision, Conceptualization, Validation, Writing – review editing, Funding acquisition.

